# The microbiome and resistome of hospital sewage during passage through the community sewer system

**DOI:** 10.1101/216242

**Authors:** Elena Buelow, Jumamurat R. Bayjanov, Rob J.L. Willems, Marc J.M. Bonten, Heike Schmitt, Willem van Schaik

## Abstract

Effluents from wastewater treatment plants (WWTPs) have been proposed to act as point sources of antibiotic-resistant bacteria (ARB) and antimicrobial resistance genes (ARGs) in the environment. Hospital sewage may contribute to the spread of ARB and ARGs as it contains the feces and urine of hospitalized patients, who are more frequently colonized with multi-drug resistant bacteria than the general population. However, whether hospital sewage noticeably contributes to the quantity and diversity of ARGs in the general sewerage system has not yet been determined.

Here, we employed culture-independent techniques, namely 16S rRNA and nanolitre-scale quantitative PCRs, to describe the role of hospital effluent as a point source of ARGs in the sewer system, through comparing microbiota composition and levels of ARGs in hospital sewage with WWTP influent, WWTP effluent and the surface water in which the effluent is released.

Compared to other sample sites, hospital sewage was richest in human-associated bacteria and contained the highest relative levels of ARGs. Yet, the abundance of ARGs was comparable in WWTPs with and without hospital wastewater, suggesting that hospitals do not contribute to the spread of ARGs in countries with a functioning sewerage system.

## Introduction

Antibiotic-producing and antibiotic-resistant bacteria (ARB) naturally and ubiquitously occur in the environment (Anukool *et al.*, 2004; Wellington *et al.*, 2013). However, human activities contribute importantly to the dissemination of resistant bacteria and resistance genes from humans and animals to the environment (Woolhouse & Ward, 2013). Effluents of wastewater treatment plants (WWTPs) may represent an important source of ARB and antimicrobial resistance genes (ARGs) in the aquatic environment (LaPara *et al.*, 2011; Wellington *et al.*, 2013; Pruden *et al.*, 2013; Rizzo *et al.*, 2013; Stalder *et al.*, 2014; Pruden, 2014; Czekalski *et al.*, 2014; Blaak *et al.*, 2014; Karkman *et al.*, 2016; Karkman *et al.*, 2017). Generally, WWTPs collect municipal wastewater, but also wastewater from industry, farms and hospitals, dependent on the size and nature of the communities connected to a single sewer system. In hospitals, up to one third of patients receive antibiotic therapy on any given day and consequently, hospitals may be important hubs for the emergence and spread of ARB and ARGs (Vlahovic-Palcevski *et al.*, 2007; Bush *et al.*, 2011; Robert *et al.*, 2012). Recent studies have highlighted that multidrug-resistant nosocomial pathogens, ARGs and genetic determinants that contribute to the mobilization and dissemination of ARGs are abundant in hospital sewage, indicating that hospital sewage may play a role in the dissemination of bacteria and genetic determinants involved in antibiotic resistance (Kummerer, 2001; Borjesson *et al.*, 2009; Stalder *et al.*, 2013; Varela *et al.*, 2013; Stalder *et al.*, 2014; Brechet *et al.*, 2014).

Whether hospital effluent contributes to the presence of ARGs in the aquatic environment is still poorly understood. To quantify the role of hospital effluent as a point source of ARGs in the sewer system, we compared the levels of ARGs in hospital sewage with WWTP influent, WWTP effluent and the surface water in which the effluent is released. In addition, we investigated the microbial composition along this sample gradient, in order to follow the fate of intestinal microbiota as sources of ARGs.

## Materials and Methods

### Sampling locations

Sampling was conducted at the main hospital wastewater pipe of the University Medical Center Utrecht (UMCU), Utrecht in the Netherlands, and at two WWTP plants. One plant (termed ‘urban WWTP’ in this manuscript) treats wastewater of approximately 290,000 inhabitants of the city of Utrecht, including the investigated hospital and two other hospitals. The other plant (‘suburban WWTP’ in Lopik, the Netherlands) neither serves a hospital nor a retirement or nursing home, and treats wastewater of a suburban community of approximately 14,000 inhabitants (Supplementary Figure 1). Both plants apply secondary treatment including nitrification and denitrification in activated sludge systems. Phosphorus removal is performed chemically in the urban WWTP, and biologically in the suburban WWTP. The hospital has approximately 1,000 beds and 8,200 employees (full-time equivalents). Additionally, some 2,500 students are enrolled at the university hospital.

**Figure 1:**
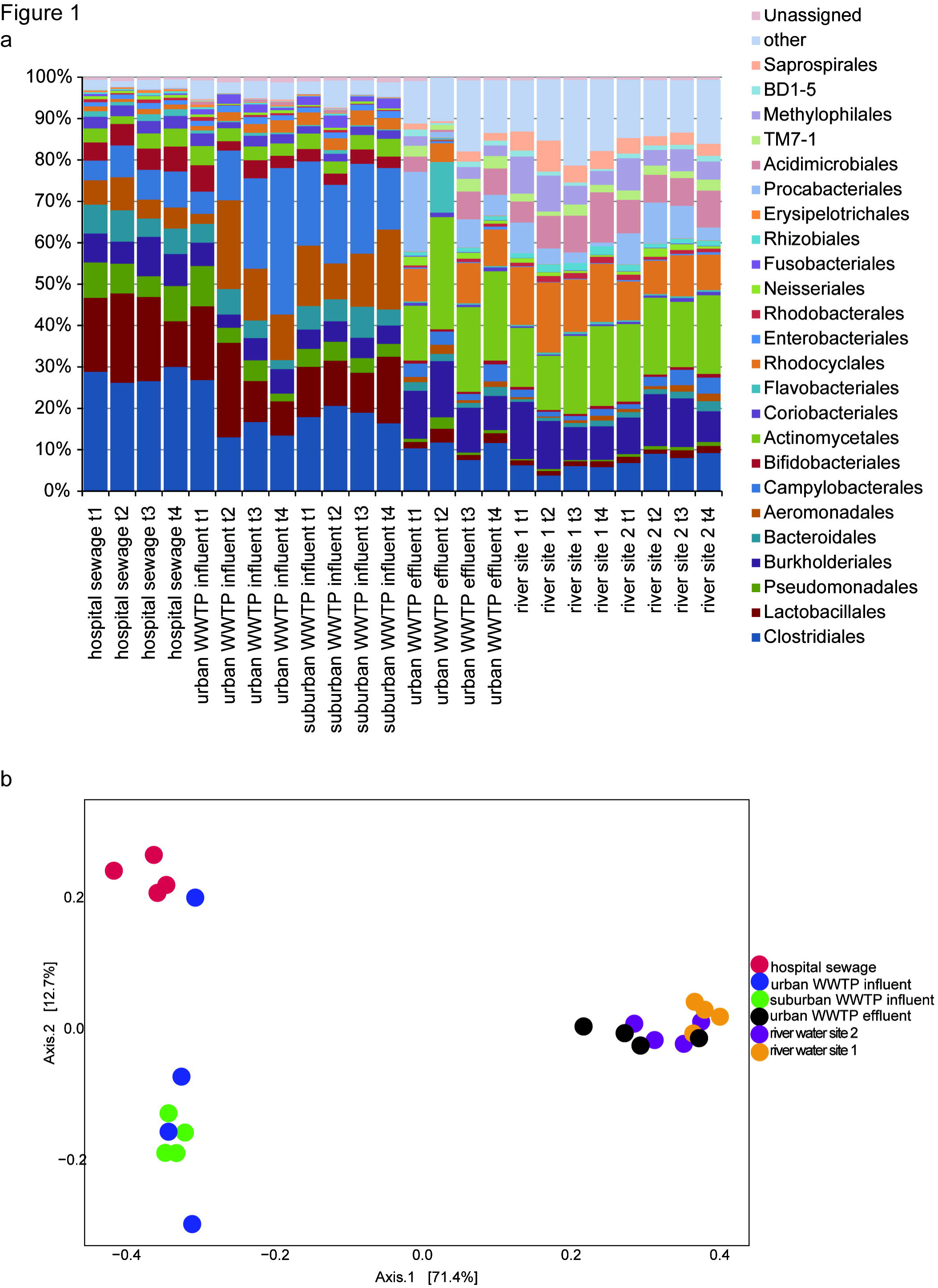
Microbiota composition of the sample locations at different time points. **A:** Relative abundance of bacteria at the order level in different samples as detected by dual indexing 16S rRNA Illumina MiSeq sequencing. The 24 most abundant bacteria at the order level for all samples are depicted, where the “other” represents percentage of the remaining taxa and “Unassigned” shows percentage of OTUs that could not be assigned to any known taxonomy. The different sampling time points are indicated as t1 (Monday 31 March 2014); t2 (Wednesday 2 April 2014); t3 (Monday 7 April 2014); t4 (Monday 14 April 2014). **B:** Principal coordinates analysis (PCoA) of microbiota composition for all different locations and time points. PCoA based on the weighted UniFrac distance depicts the differences in microbiota compositions.

### Sampling and DNA isolation

Samples were taken during a period of 2.5 weeks in spring on four days (Monday 31 March 2014; Wednesday 2 April 2014; Monday 7 April 2014 and Monday 14 April 2014). Cumulative precipitation in the three days preceding each sampling date amounted to maximally 15 mm. The daily flows amount to 74,800 ± 5,900 m^3^ for the urban WWTP, and 3,390 ± 380 m^3^ for the suburban WWTP during the four sampling days. The flows of the academic hospital amount to approximately 216,000 m^3^ on a yearly basis, i.e. on average 590 m^3^ per day (0.8% of the influent of the urban WWTP). Exact quantification of the flows of the academic hospital is not possible as the daily flows are not regularly registered. Flow-proportional sampling (over 24 hours) was used for sampling hospital wastewater and WWTP influent and effluent. Samples were kept at 4°C during the flow-proportional sampling. For the surface water samples, grab samples (5 L) were taken at 50 cm downstream of the two effluent pipes of the urban WWTP discharging into a local river at a depth of 20 cm. Samples were transported to the laboratory at 4°C and samples were processed the same day. Bacterial cell pellets and debris of sewage and surface water samples were collected by means of centrifugation at 14000 *g* for 25 minutes. All pellets were resuspended in phosphate buffered saline (PBS; 138 mM NaCl, 2.7 mM KCl, 140 mM Na_2_HPO_4_, 1.8 mM KH_2_PO_4_, adjusted to pH 7.4 with HCl) with 20% glycerol and stored at −80 C° until DNA extraction. Metagenomic DNA was extracted from 200 µl of frozen samples as described previously (Godon *et al.*, 1997).

### 16S rRNA sequencing and sequence data pre-processing

16S rRNA sequencing was performed on the Illumina MiSeq sequencing platform (San Diego, CA). A dual-indexing approach for multiplex 16S rRNA sequencing targeting the V3-V4 hypervariable region of the 16S rRNA gene was employed as described by (Fadrosh *et al.*, 2014), using the 300 bp paired-end protocol to sequence a pool of 24 samples. Untrimmed paired-end reads were assembled using the FLASH assembler, which performs error correction during the assembly process (Magoc & Salzberg, 2011). After removal of the barcodes, heterogeneity spacers, and primer sequences, there were a total of 1.4 million joined reads with a median length of 424 bases and a median number of 57860 joined reads per sample.

### 16S rRNA sequence data analysis

Joined reads were further analyzed using the QIIME microbial community analysis pipeline (version 1.8.0) (Caporaso, Kuczynski *et al.*, 2010). Joined reads with a minimum of 97% similarity were assigned into operational taxonomic units (OTUs) using QIIME’s open-reference OTU calling workflow. This workflow was used with the “-m usearch61” option, which uses the USEARCH algorithm (Edgar, 2010) for OTU picking and UCHIME for chimeric sequence detection (Edgar *et al.*, 2011). Taxonomic ranks for OTUs were assigned using the Greengenes database (version 13.8) (McDonald *et al.*, 2012) with the default parameters of the script pick_open_reference_otus.py. A representative sequence of each OTU was aligned to the Greengenes core reference database (DeSantis *et al.*, 2006) using the PyNAST aligner (version 1.2.2) (Caporaso, Bittinger *et al.*, 2010). Highly variable parts of alignments were removed using the filter_alignment.py script, which is part of the pick_open_reference_otus.py workflow. Subsequently, filtered alignment results were used to create an approximate maximum-likelihood phylogenetic tree using FastTree (version 2.1.3) (Price *et al.*, 2010). For more accurate taxa diversity distribution (Bokulich *et al.*, 2013), OTUs to which less than 0.005% of the total number of assembled reads were mapped, were discarded using the filter_otus_from_otu_table.py script with the parameter “--min_count_fraction 0.00005”. The filtered OTU table and generated phylogenetic tree were used to assess within-sample (alpha) and between sample (beta) diversities.

Alpha- and beta-diversity of samples were assessed using QIIME’s core_diversity_analyses.py workflow. For rarefaction analysis the subsampling depth threshold of 20681 was used, which was the minimum number of reads assigned to a sample. The UniFrac distance was used as input to calculate the Chao1 index as a measure of beta-diversity of the samples (C. Lozupone & Knight, 2005). In addition to alpha- and beta-diversity analysis and visualizations, this workflow also incorporates principal coordinates analysis and visualization of sample compositions using Emperor (Vazquez-Baeza *et al.*, 2013). Differences in the abundance of taxa are shown as averages over the four time points ± standard deviation resulting in six different comparisons between the different samples. The non-parametric Mann-Whitney test was used to test for statistical significance.

### High-throughput qPCR

Real-Time PCR analysis was performed using the 96.96 BioMark™ Dynamic Array for Real-Time PCR (Fluidigm Corporation, San Francisco, CA, U.S.A), according to the manufacturer’s instructions, with the exception that the annealing temperature in the PCR was lowered to 56°C.

Other technical details of the nanolitre-scale quantitative PCRs to quantify levels of genes that confer resistance to antimicrobials (antibiotics and disinfectants, specifically quaternary ammonium compounds (QACs)) were described previously (Buelow *et al.*, 2017), with some modifications in the collection of primers. Primers that were negative for all samples tested in the previous study, or gave unspecific amplification were redesigned or replaced. The primer sequences and their targets are provided in the supplementary data, updated and novel primers and targets are highlighted with a grey background (Supplementary Table 1).

### Calculation of normalized abundance and cumulative abundance

Normalized abundance of resistance genes was calculated relative to the abundance of the 16S rRNA gene (CT_ARG_ – CT_16S rRNA_) resulting in a log2-transformed estimate of ARG abundance. Cumulative abundance was calculated based on the sum of the non-log2 transformed values (2^(-(CT_ARG_ – CT_16S rRNA_)) of all genes within a resistance gene family. The differences in cumulative abundance over all time points are shown as an averaged fold-change ± standard deviation. The non-parametric Mann-Whitney test was used to test for significance; p values were corrected for multiple testing by the Benjamin-Hochberg procedure (Benjamini & Hochberg, 1995) with a false discovery rate of 0.05.

## Results

### Composition of the microbiota of hospital sewage, WWTP influent, WWTP effluent and river water

The composition of the microbiota in hospital sewage, urban and suburban WWTP influents, the effluent of the urban WWTP and the surface water in which the effluent was released was determined by multiplexed 16S rRNA sequencing on the Illumina MiSeq platform (Figure 1A and Supplementary Table 2). At all sample sites, the microbiota consisted of a complex consortium of bacteria from different orders with the microbiota being most diverse in the river samples and least diverse in hospital sewage (Supplementary Figure 2). Hospital sewage contained relatively high levels (39.4 ± 2.5% standard deviation, of the total microbiota) of anaerobic bacteria (Bifidobacteriales, Bacteroidales and Clostridiales) that are likely to originate from the human gut (Rajilic-Stojanovic & de Vos, 2014). These orders were less abundant in WWTP influent (25.8 ± 8.4%) and suburban WWTP influent (27 ± 3%; *p* < 0.05) compared to hospital sewage. Compared to the WWTP influent, abundance of Bifidobacteriales, Bacteroidales and Clostridiales was significantly (*p* < 0.05) lower in WWTP effluent (12.8 ± 2.5%) and river water (7.0 ± 1.3% for site 1 and 10.6 ± 1.6% for site 2). In contrast, bacteria that are associated with activated sludge (such as the Actinomycetales, Rhodocyclales, and Burkholderiales (Zhang *et al.*, 2012)) became more prominent during passage through the sewer system and WWTP (Figure 1A and Supplementary Table 2). Principal coordinates analysis (PCoA) showed a clear distinction between the samples that were isolated prior to treatment in the WWTP and the samples of WWTP effluent and river water (Figure 1B). The three most abundant bacterial taxa detected in the hospital sewage were the genera *Streptococcus* (9.0%) and *Arcobacter* (6.9%) and the family Ruminococcaceae (6.3%). Both raw sewage influents (urban WWTP influent, suburban WWTP influent) clustered together and in both sites, the same three bacterial taxa were most abundant (*Arcobacter*: 17.9% in urban WWTP influent; 17.5% in suburban WWTP influent; Aeromonadaceae: 11.2% and 12.4% respectively; Carnobacteriaceae, 9.4% and 8.3% respectively). Variation of the microbiota composition between the different samples dates at the six sites was limited; with the urban WWTP influent and effluent exhibiting the most pronounced fluctuations (Figure 1b). The urban WWTP effluent samples were very similar to the surface water samples that were collected close to the effluent release pipes. Urban WWTP effluent shared the same three most common OTUs with one of the surface water samples (Actinomycetales, 15.4% in urban WWTP effluent and 9.7% in river site 2; Procabacteriaceae, 8.1% and 7.1% respectively; Comamonadaceae, 7.6% and 7.7% respectively). The surface water sample collected at the other release pipe (river site 1) was slightly different and is defined by the following three most abundant OTUs: Comamonadaceae, 7.5%, Intrasporangiaceae, 6.1% and *Candidatus Microthrix*, 6.1%.

**Figure 2:**
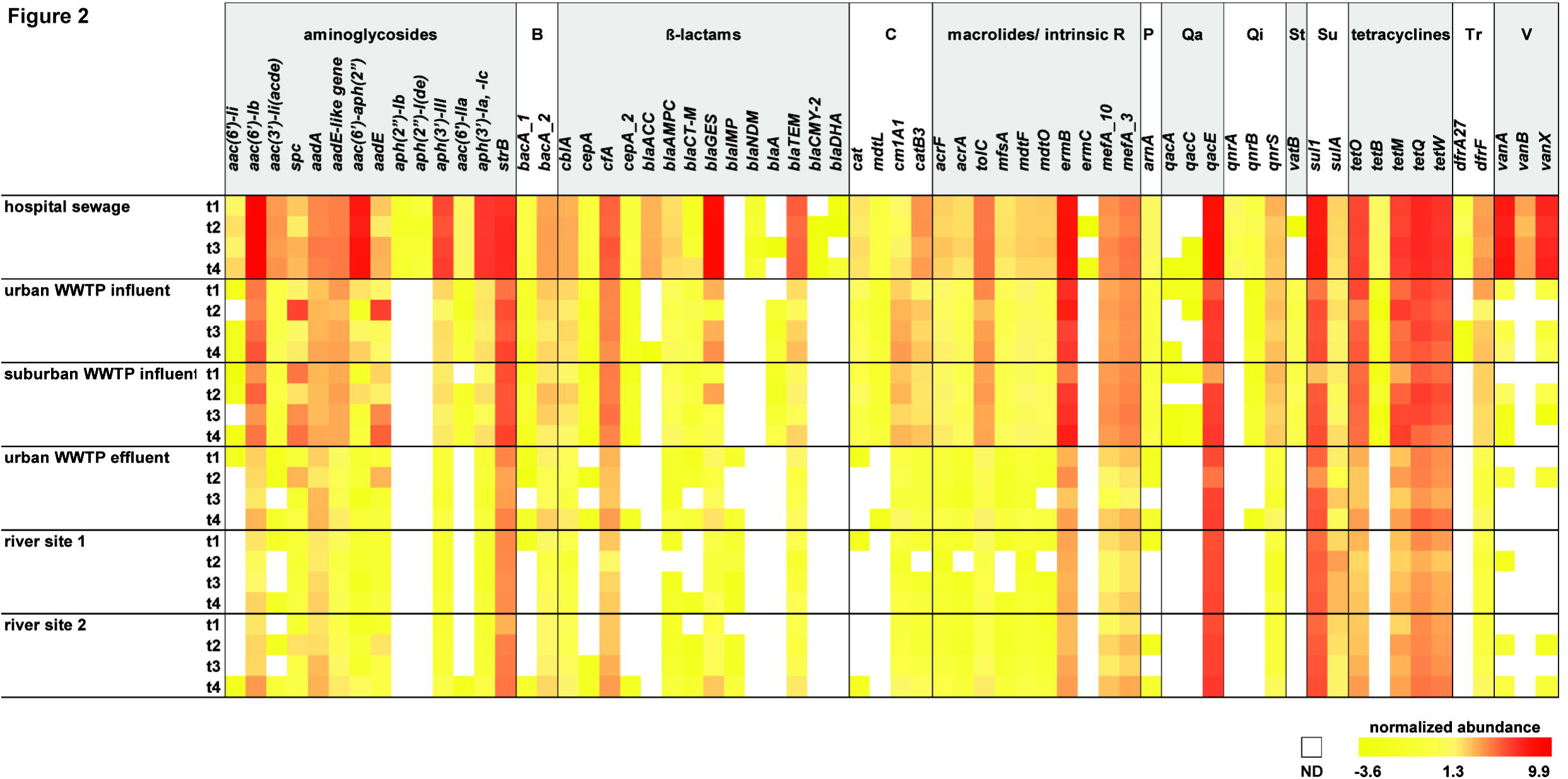
Abundance levels of ARGs in hospital, WWTP influent, WWTP effluent and river water. 16S rRNA - normalized abundance of ARGs detected in all samples. ARGs are grouped according to resistance gene families (aminoglycosides; B, bacitracin, β-lactams; C, chloramphenicols; macrolide / intrinsic resistance; P, polymyxins; Pu, puromycins; Qa, QAC resistance genes; Qi, quinolones; St, streptogramins; Su, sulphonamides; tetracyclines; Tr, trimethoprim; V, vancomycin). The colour scale ranges from bright red (most abundant) to bright yellow (least abundant). White blocks indicate that a resistance gene was not detected. The different sampling time points are indicated as t1 (Monday 31 March 2014); t2 (Wednesday 2 April 2014); t3 (Monday, 7 April 2014); t4 (Monday, 14 April 2014).

### Resistome composition of hospital sewage compared to receiving urban sewage

A total of 67 ARGs were detected in the different samples, conferring resistance to 13 classes of antimicrobials. The levels of ARGs were calculated as a normalized abundance relative to levels of 16S rRNA, which provides an indication of the relative levels of ARGs within the bacterial population in each sample (Figure 2, Supplementary Table 3). Hospital sewage was found to be enriched in ARGs, compared to the other samples. The normalized abundance of 11 out of 13 classes of ARGs was significantly (*p*<0.05) higher in hospital sewage than in the urban WWTP influent, particularly so for aminoglycoside (12.0 ± 5.0-fold higher in hospital sewage), β-lactam (15.4 ± 3.6-fold higher in hospital sewage) and vancomycin resistance genes (175 ± 14-fold higher in hospital sewage, based on the three days when vancomycin resistance genes could be detected in the WWTP influent). Only one class of resistance genes (conferring resistance to streptogramins) was significantly less abundant (*p*<0.05) in hospital sewage than in WWTP influent. The combined levels of chloramphenicol and quinolone resistance genes were not different between the sites. Seven ARGs (two aminoglycoside resistance genes, *aph(2”)-Ib* and *aph(2”)-I(de),* the quinolone resistance gene *qnrA,* the, erythromycin resistance gene *ermC,* the vancomycin resistance gene *vanB*, the AmpC-type β-lactamases *bla*_DHA-1_ and *bla*_CMY-2_ and the carbapenemase *bla*_NDM_) were only detected in hospital sewage (Figure 2). The abundance of ARGs in the urban WWTP influent, which receives sewage from the sampled hospital and two additional hospitals in the same city, and the suburban WWTP influent is comparable (Figure 2 Supplementary Table 3). For nine classes of antibiotics, (aminoglycosides, β-lactams, chloramphenicols, macrolides, polymyxins, puromycins, trimpethoprim, quinolones, and tetracyclines), the levels of ARGs in the urban WWTP effluent were significantly (*p* <0.05) lower than in the WWTP influent (ranging between a 7.6 ± 2.2-fold reduction for macrolide and intrinsic (efflux) resistance genes to a 2.8 ± 0.9-fold reduction for β-lactam resistance genes), with the remaining classes of ARGs not changing significantly in abundance (Figure 2 and Supplementary Table 3). The levels of ARGs in WWTP effluent were comparable to the levels of ARGs in river water (Figure 2, Supplementary Table 3).

## Discussion

Our study demonstrates the considerable efflux of ARGs from hospitals through sewage. However, the influents of the urban and suburban WWTPs studied here show very similar levels of ARGs, even though the urban WWTP receives sewage from a variety of sources including three hospitals, while the sub-urban WWTP does not have a hospital in its catchment area. This reflects the relatively limited effect of hospital sewage on the level of ARGs in WWTP influent and the low contribution of hospital sewage (an estimated 0.8%) to the total volume of wastewater treated in the urban WWTP that we investigated. Our study further demonstrates the capacity of WWTPs to reduce the ultimate amount of ARGs present in urban and suburban influent before entering the river systems.

Effluents from WWTPs are thought to contribute to the dissemination of pollutants, multi-drug resistant bacteria and resistance genes in the environment (Rizzo *et al.*, 2013; Wellington *et al.*, 2013; Karkman *et al.*, 2017). Particularly high amounts of ARB have previously been reported in hospital sewage (Diwan *et al.*, 2010; Wellington *et al.*, 2013; Harris *et al.*, 2013; Harris *et al.*, 2014; Berendonk *et al.*, 2015). Large amounts of antibiotics and QACs are used in hospitals and these may promote the establishment of ARB and selection of ARGs in patients and hospital wastewaters (Stalder *et al.*, 2013; Stalder *et al.*, 2014; Varela *et al.*, 2014). Here we show that the relative abundance of a broad range of ARGs conferring resistance to 11 classes of antimicrobials is significantly higher in hospital sewage compared to urban and suburban WWTP sewage. In particular, genes conferring resistance to aminoglycosides, β-lactams and vancomycin are enriched in hospital sewage, presumably due to the frequent use of these classes of antibiotics in the hospital (Chandy *et al.*, 2014).

The most abundant bacterial taxa detected in the hospital sewage are different from those found in the urban and suburban WWTP influent, which are dominated by bacterial families (*Arcobacter*; Aeromonadaceae; Carnobacteriaceae) that are commonly found in the sewer microbiota (Moreno *et al.*, 2003; Vandewalle *et al.*, 2012; Shanks *et al.*, 2013). Compared to the WWTP influent samples, several members of the human gut microbiota are significantly more abundant in hospital sewage, most probably due to the close proximity of the sampling location to the hospital sanitation systems. These human-associated taxa include the genus *Streptococcus*, of which many species interact with humans either as commensals or pathogens (Kalia *et al.*, 2001), and the Ruminococcaceae, which are one of the most prevalent bacterial families in the human gut (Arumugam *et al.*, 2011; C. A. Lozupone *et al.*, 2012). These human-associated bacteria appear to be ill-suited for surviving the complex and, at least partially oxygenated, sewage environment and progressively decrease in abundance, leading to lower levels of human gut-associated bacteria in the urban WWTP influent. Because most ARGs from the human microbiota appear to be carried by non-pathogenic commensal bacteria (Sommer *et al.*, 2009; Buelow *et al.*, 2014), a general loss of human commensal bacteria in the sewer system (Pehrsson *et al.*, 2016) may contribute to a decrease in the abundance of ARGs during the passage of wastewater through the sewer system.

The reduction of ARGs shown in urban WWTP effluent compared to WWTP influent may be explained by the further significant reduction of the relative abundance of human-associated bacterial taxa detected in urban WWTP effluent. The continuous reduction of these bacterial taxa could be mediated by their removal through sorption to activated sludge, by replacement with the bacteria that populate activated sludge, and/or by predation of protozoa during wastewater treatment (Wen *et al.*, 2009; Calero-Caceres *et al.*, 2014). Interestingly, the presence of Procabacteriales in WWTP effluent and river water (Figure 1A), may point towards a relatively high abundance of protists in these samples, as these bacteria are intracellular symbionts or pathogens of amoeba (Horn *et al.*, 2002; Greub & Raoult, 2004).

The reduction of the abundance of ARGs from hospital sewage to WWTP effluent highlights the importance of wastewater treatment in reducing the discharge of ARGs originating from human sources into the environment. With respect to the abundance of ARG relative to 16S rRNA, it has been debated whether sewage treatment could selectively affect the percentage of resistant bacteria within a given species, or within the total community (Rizzo *et al.*, 2013; Laht *et al.*, 2014; Alexander *et al.*, 2015). Here, and in line with (Karkman *et al.*, 2016), we observed that wastewater treatment led to a decrease in the relative abundance of ARGs. A clear decrease of ARGs in absolute terms (i.e. in gene copies per volume of water) along the sewage treatment has been shown in a range of publications (Auerbach *et al.*, 2007; Chen & Zhang, 2013; Laht *et al.*, 2014; Czekalski *et al.*, 2014).

Sampling for this study was limited to one single season, but was repeated four consecutive days in dry weather conditions using mostly flow-proportional sampling as recommended by Ort *et al.* (Ort *et al.*, 2010). Microbiota and resistome profiling of our samples showed limited variation between the four sampling days for each sample, hence allowing for analysis of the treatment efficacy on the removal of ARGs relative to 16S rRNA in these particular WWTPs.

Advanced water treatment methods have been proposed as a selective measure for hospital wastewater, specifically to decrease pharmaceuticals and the release of pathogens by hospitals (Lienert *et al.*, 2011). For the investigated municipal WWTP, hospital wastewater seems to play a limited role for the level of resistance genes in the influent and effluent. Our findings suggest that -in the presence of operational WWTPs-hospital-specific sewage treatment will not lead to a substantial further reduction of the release of ARGs into effluent.

## References

Alexander J, Bollmann A, Seitz W & Schwartz T (2015) Microbiological characterization of aquatic microbiomes targeting taxonomical marker genes and antibiotic resistance genes of opportunistic bacteria Sci Total Environ 512-513: 316–325.

Anukool U, Gaze WH & Wellington EM (2004) In situ monitoring of streptothricin production by Streptomyces rochei F20 in soil and rhizosphere Appl Environ Microbiol 70: 5222–5228.

Arumugam M, Raes J, Pelletier E et al. (2011) Enterotypes of the human gut microbiome Nature 473: 174–180.

Auerbach EA, Seyfried EE & McMahon KD (2007) Tetracycline resistance genes in activated sludge wastewater treatment plants Water Res 41: 1143–1151.

Benjamini Y & Hochberg Y (1995) Controlling the false discovery rate: a practical and powerful approach to multiple testing. Journal of the Royal Statistical Society Series B (Methodological) Vol. 57, No. 1: pp. 289–300.

Berendonk TU, Manaia CM, Merlin C et al. (2015) Tackling antibiotic resistance: the environmental framework Nat Rev Microbiol 13: 310–317.

Blaak H, de Kruijf P, Hamidjaja RA, van Hoek AH, de Roda Husman AM & Schets FM (2014) Prevalence and characteristics of ESBL-producing E. coli in Dutch recreational waters influenced by wastewater treatment plants Vet Microbiol 171: 448–459.

Bokulich NA, Subramanian S, Faith JJ, Gevers D, Gordon JI, Knight R, Mills DA & Caporaso JG (2013) Quality-filtering vastly improves diversity estimates from Illumina amplicon sequencing Nat Methods 10: 57–59.

Borjesson S, Dienues O, Jarnheimer PA, Olsen B, Matussek A & Lindgren PE (2009) Quantification of genes encoding resistance to aminoglycosides, beta-lactams and tetracyclines in wastewater environments by real-time PCR Int J Environ Health Res 19:219–230.

Brechet C, Plantin J, Sauget M, Thouverez M, Talon D, Cholley P, Guyeux C, Hocquet D & Bertrand X (2014) Wastewater treatment plants release large amounts of extended-spectrum beta-lactamase-producing Escherichia coli into the environment Clin Infect Dis 58: 1658–1665.

Buelow E, Bello Gonzalez TDJ, Fuentes S et al. (2017) Comparative gut microbiota and resistome profiling of intensive care patients receiving selective digestive tract decontamination and healthy subjects Microbiome 5: 88-017-0309-z.

Buelow E, Gonzalez TB, Versluis D et al. (2014) Effects of selective digestive decontamination (SDD) on the gut resistome J Antimicrob Chemother. 69: 2215–23

Bush K, Courvalin P, Dantas G et al. (2011) Tackling antibiotic resistance Nat Rev Microbiol 9: 894–896.

Calero-Caceres W, Melgarejo A, Colomer-Lluch M, Stoll C, Lucena F, Jofre J & Muniesa M (2014) Sludge as a potential important source of antibiotic resistance genes in both the bacterial and bacteriophage fractions Environ Sci Technol 48: 7602–7611.

Caporaso JG, Bittinger K, Bushman FD, DeSantis TZ, Andersen GL & Knight R (2010) PyNAST: a flexible tool for aligning sequences to a template alignment Bioinformatics 26: 266–267.

Caporaso JG, Kuczynski J, Stombaugh J et al. (2010) QIIME allows analysis of high-throughput community sequencing data Nat Methods 7: 335–336.

Chandy SJ, Naik GS, Charles R, Jeyaseelan V, Naumova EN, Thomas K & Lundborg CS (2014) The impact of policy guidelines on hospital antibiotic use over a decade: a segmented time series analysis PLoS One 9: e92206.

Chen H & Zhang M (2013) Occurrence and removal of antibiotic resistance genes in municipal wastewater and rural domestic sewage treatment systems in eastern China Environ Int 55: 9–14.

Czekalski N, Gascon Diez E & Burgmann H (2014) Wastewater as a point source of antibiotic-resistance genes in the sediment of a freshwater lake ISME J 8: 1381–1390.

DeSantis TZ, Hugenholtz P, Larsen N, Rojas M, Brodie EL, Keller K, Huber T, Dalevi D, Hu P & Andersen GL (2006) Greengenes, a chimera-checked 16S rRNA gene database and workbench compatible with ARB Appl Environ Microbiol 72: 5069–5072.

Diwan V, Tamhankar AJ, Khandal RK, Sen S, Aggarwal M, Marothi Y, Iyer RV, Sundblad-Tonderski K & Stalsby-Lundborg C (2010) Antibiotics and antibiotic-resistant bacteria in waters associated with a hospital in Ujjain, India BMC Public Health 10: 414-2458-10-414.

Edgar RC (2010) Search and clustering orders of magnitude faster than BLAST Bioinformatics 26: 2460–2461.

Edgar RC, Haas BJ, Clemente JC, Quince C & Knight R (2011) UCHIME improves sensitivity and speed of chimera detection Bioinformatics 27: 2194–2200.

Fadrosh DW, Ma B, Gajer P, Sengamalay N, Ott S, Brotman RM & Ravel J (2014) An improved dual-indexing approach for multiplexed 16S rRNA gene sequencing on the Illumina MiSeq platform Microbiome 2: 6-2618-2-6.

Godon JJ, Zumstein E, Dabert P, Habouzit F & Moletta R (1997) Molecular microbial diversity of an anaerobic digestor as determined by small-subunit rDNA sequence analysis Appl Environ Microbiol 63: 2802–2813.

Greub G & Raoult D (2004) Microorganisms resistant to free-living amoebae Clin Microbiol Rev 17: 413–433.

Harris S, Morris C, Morris D, Cormican M & Cummins E (2014) Antimicrobial resistant *Escherichia coli* in the municipal wastewater system: effect of hospital effluent and environmental fate Sci Total Environ 468-469: 1078–1085.

Harris S, Morris C, Morris D, Cormican M & Cummins E (2013) The effect of hospital effluent on antimicrobial resistant E. coli within a municipal wastewater system Environ Sci Process Impacts 15: 617–622.

Horn M, Fritsche TR, Linner T, Gautom RK, Harzenetter MD & Wagner M (2002) Obligate bacterial endosymbionts of Acanthamoeba spp. related to the beta-Proteobacteria: proposal of ’Candidatus Procabacter acanthamoebae’ gen. nov., sp. nov Int J Syst Evol Microbiol 52: 599–605.

Kalia A, Enright MC, Spratt BG & Bessen DE (2001) Directional gene movement from human-pathogenic to commensal-like streptococci Infect Immun 69: 4858–4869.

Karkman A, Do TT, Walsh F & Virta MPJ (2017) Antibiotic-Resistance Genes in Waste Water Trends Microbiol. pii: S0966-842X(17)30210-X.

Karkman A, Johnson TA, Lyra C, Stedtfeld RD, Tamminen M, Tiedje JM & Virta M (2016) High-throughput quantification of antibiotic resistance genes from an urban wastewater treatment plant FEMS Microbiol Ecol 92: 10.1093/femsec/fiw014. Epub 2016 Jan 31.

Kummerer K (2001) Drugs in the environment: emission of drugs, diagnostic aids and disinfectants into wastewater by hospitals in relation to other sources--a review Chemosphere 45: 957–969.

Laht M, Karkman A, Voolaid V, Ritz C, Tenson T, Virta M & Kisand V (2014) Abundances of tetracycline, sulphonamide and beta-lactam antibiotic resistance genes in conventional wastewater treatment plants (WWTPs) with different waste load PLoS One 9: e103705.

LaPara TM, Burch TR, McNamara PJ, Tan DT, Yan M & Eichmiller JJ (2011) Tertiary-treated municipal wastewater is a significant point source of antibiotic resistance genes into Duluth-Superior Harbor Environ Sci Technol 45: 9543–9549.

Lienert J, Koller M, Konrad J, McArdell CS & Schuwirth N (2011) Multiple-criteria decision analysis reveals high stakeholder preference to remove pharmaceuticals from hospital wastewater Environ Sci Technol 45: 3848–3857.

Lozupone C & Knight R (2005) UniFrac: a new phylogenetic method for comparing microbial communities Appl Environ Microbiol 71: 8228–8235.

Lozupone CA, Stombaugh JI, Gordon JI, Jansson JK & Knight R (2012) Diversity, stability and resilience of the human gut microbiota Nature 489: 220–230.

Magoc T & Salzberg SL (2011) FLASH: fast length adjustment of short reads to improve genome assemblies Bioinformatics 27: 2957–2963.

McDonald D, Price MN, Goodrich J, Nawrocki EP, DeSantis TZ, Probst A, Andersen GL, Knight R & Hugenholtz P (2012) An improved Greengenes taxonomy with explicit ranks for ecological and evolutionary analyses of bacteria and archaea ISME J 6: 610–618.

Moreno Y, Botella S, Alonso JL, Ferrus MA, Hernandez M & Hernandez J (2003) Specific detection of Arcobacter and Campylobacter strains in water and sewage by PCR and fluorescent in situ hybridization Appl Environ Microbiol 69: 1181–1186.

Ort C, Lawrence MG, Rieckermann J & Joss A (2010) Sampling for pharmaceuticals and personal care products (PPCPs) and illicit drugs in wastewater systems: are your conclusions valid? A critical review Environ Sci Technol 44: 6024–6035.

Pehrsson EC, Tsukayama P, Patel S et al. (2016) Interconnected microbiomes and resistomes in low-income human habitats Nature 533: 212–216.

Price MN, Dehal PS & Arkin AP (2010) FastTree 2--approximately maximum-likelihood trees for large alignments PLoS One 5: e9490.

Pruden A (2014) Balancing water sustainability and public health goals in the face of growing concerns about antibiotic resistance Environ Sci Technol 48: 5–14.

Pruden A, Larsson DG, Amezquita A et al. (2013) Management options for reducing the release of antibiotics and antibiotic resistance genes to the environment Environ Health Perspect 121: 878–885.

Rajilic-Stojanovic M & de Vos WM (2014) The first 1000 cultured species of the human gastrointestinal microbiota FEMS Microbiol Rev 38: 996–1047.

Rizzo L, Manaia C, Merlin C, Schwartz T, Dagot C, Ploy MC, Michael I & Fatta-Kassinos D (2013) Urban wastewater treatment plants as hotspots for antibiotic resistant bacteria and genes spread into the environment: a review Sci Total Environ 447: 345–360.

Robert J, Pean Y, Varon E et al. (2012) Point prevalence survey of antibiotic use in French hospitals in 2009 J Antimicrob Chemother 67: 1020–1026.

Shanks OC, Newton RJ, Kelty CA, Huse SM, Sogin ML & McLellan SL (2013) Comparison of the microbial community structures of untreated wastewaters from different geographic locales Appl Environ Microbiol 79: 2906–2913.

Sommer MO, Dantas G & Church GM (2009) Functional characterization of the antibiotic resistance reservoir in the human microflora Science 325: 1128–1131.

Stalder T, Barraud O, Jove T, Casellas M, Gaschet M, Dagot C & Ploy MC (2014) Quantitative and qualitative impact of hospital effluent on dissemination of the integron pool ISME J 8: 768–777.

Stalder T, Alrhmoun M, Louvet JN, Casellas M, Maftah C, Carrion C, Pons MN, Pahl O, Ploy MC & Dagot C (2013) Dynamic assessment of the floc morphology, bacterial diversity, and integron content of an activated sludge reactor processing hospital effluent Environ Sci Technol 47: 7909–7917.

Vandewalle JL, Goetz GW, Huse SM, Morrison HG, Sogin ML, Hoffmann RG, Yan K & McLellan SL (2012) Acinetobacter, Aeromonas and Trichococcus populations dominate the microbial community within urban sewer infrastructure Environ Microbiol 14: 2538–2552.

Varela AR, Andre S, Nunes OC & Manaia CM (2014) Insights into the relationship between antimicrobial residues and bacterial populations in a hospital-urban wastewater treatment plant system Water Res 54: 327–336.

Varela AR, Ferro G, Vredenburg J, Yanik M, Vieira L, Rizzo L, Lameiras C & Manaia CM (2013) Vancomycin resistant enterococci: from the hospital effluent to the urban wastewater treatment plant Sci Total Environ 450-451: 155–161.

Vazquez-Baeza Y, Pirrung M, Gonzalez A & Knight R (2013) EMPeror: a tool for visualizing high-throughput microbial community data Gigascience 2: 16-217X-2-16.

Vlahovic-Palcevski V, Dumpis U, Mitt P, Gulbinovic J, Struwe J, Palcevski G, Stimac D, Lagergren A & Bergman U (2007) Benchmarking antimicrobial drug use at university hospitals in five European countries Clin Microbiol Infect 13: 277–283.

Wellington EM, Boxall AB, Cross P et al. (2013) The role of the natural environment in the emergence of antibiotic resistance in gram-negative bacteria Lancet Infect Dis 13: 155–165.

Wen Q, Tutuka C, Keegan A & Jin B (2009) Fate of pathogenic microorganisms and indicators in secondary activated sludge wastewater treatment plants J Environ Manage 90: 1442–1447.

Woolhouse ME & Ward MJ (2013) Microbiology. Sources of antimicrobial resistance Science 341: 1460–1461.

Zhang T, Shao MF & Ye L (2012) 454 Pyrosequencing Reveals Bacterial Diversity of Activated Sludge from 14 Sewage Treatment Plants ISME J 6: 1137–1147.

